# Differential Transcription of Escherichia Coli K-12 Genes Under Hypomagnetic and Geomagnetic Conditions

**DOI:** 10.64898/2026.08.03.742565

**Authors:** Michael Montague, Morgan L. Sosa, Clarice D. Aiello

## Abstract

The transcriptomic response of *Escherichia coli* to hypomagnetic versus geomagnetic exposure was examined using RNA sequencing across seven pairwise comparisons of differently conditioned cultures. Eighty-one genes were reproducibly differentially expressed, meeting significance and fold-change criteria in at least four of seven comparisons. The direction of differential expression tracked the 90-minute magnetic field exposure condition rather than the magnetic field conditions to which the seed cultures were exposed, indicating that the response is driven by acute field exposure rather than culture history. The affected genes point to a distinct metabolic state layered on top of lag phase, marked by atypical iron stress, downregulation of the flagellar regulon, and a phosphotransferase-mediated carbon-metabolism program favoring rapid energy acquisition over growth.

## 1. Introduction

Previously, we reported that hypomagnetic conditions of approximately 19 nT reproducibly lengthened the lag phase of the *Escherichia coli* growth curve in rich media by 45 minutes (almost three generation lengths) [1]. Here, we present transcriptomic surveys of differentially expressed genes during that altered lag phase. To situate these results, we first review three decades of studies documenting changes in *E. coli* gene expression in response to magnetic fields, which fall into two broad classes.

The first class of studies demonstrated limited magnetic field effects in the expression of only a few genes, even if indicative of larger transcriptomic programs. For example, Cairo *et al.* observed that sigma factor *σ*^32^ mRNA levels are enhanced following a 15-minute exposure at frequency 60 Hz and strength 1.1 mT [2]. This has broader gene-regulatory implications, as *σ*^32^ is a stress-response transcriptional modulator. This observation seems to be consistent with Potenza *et al.*’s findings upon exposure of *E. coli* to a 300 mT static magnetic field, which showed enhanced differential expression, during stationary phase, of the putative transposase TnpB [3]. This is in keeping with Cairo *et al.*’s results, as transposon activation is associated with stress response in many organisms including *E. coli* [4]. Moreover, Tsuchiya *et al.* investigated *E. coli* growing in a chamber maintaining a static field between 5.2–6.1 T. They linked higher cell titer at stationary phase under such strong magnetic fields with the activity of another sigma factor, *σ^S^*, which is encoded by the *rpoS* gene and associated with stationary phase gene expression [5].

The second class of studies investigating gene regulation with regard to magnetic field exposure in *E. coli* looks at gene expression systematically. Today, we would call this approach transcriptomics or proteomics, but the first such study predates those terms. E.M. Goodman *et al.*, in 1994, used two-dimensional protein gels to gain a proteomic-level survey of differentially expressed genes in *E. coli* exposed to a pulsed 1.5 mT magnetic field for 60 minutes [6]. These included the upregulation of NusA, a transcription elongation factor, suggesting a globally altered transcription activity. While this was a whole-proteome analysis, the 2-dimensional gel methodology and the early, pre-genomic framing makes Goodman *et al.*’s study more representative of the first limited class of early *E. coli* magnetic field gene expression work.

The first truly quantitative, sequence-based transcriptomic study was Huwiler *et al.*’s in 2012, which used an Affymetrix whole-genome microarray to measure RNA expression across all 4,358 genes and 714 intergenic regions of the *E. coli* genome. With this tool, no statistical differential expression of *E. coli* RNA products was found as a consequence of exposure to a magnetic field at frequency 50 Hz and strength 1 mT, for multiple exposure times ranging from 2.5 to 15 hours [7]. Conversely, Li *et al.* performed transcriptional analysis of *E. coli* exposed to a 250 mT static magnetic field and found a number of differentially expressed genes primarily involved in carbon source utilization. This differential gene expression was correlated to a reduced diameter of colonies (equivalent to slower or delayed growth in liquid media). Consistently, their addition of glycolate or glyoxylate to the culture media successfully restored the bacterial phenotype under the mentioned magnetic field, and knockout mutants lacking glycolate oxidase were found to no longer exhibit smaller colonies.

It should be pointed out that the results of these past studies cannot be compared directly to one another since they span a wide variety of magnetic field strengths and frequencies. Similarly, different *E. coli* strains were employed and grown in different conditions and media formulations, for different periods of time, and to different titres. Nonetheless, a broad trend of magnetic fields causing stress responses was seen across most, but not all, of the studies. This is in keeping with Li *et al.*’s more recent 2025 report, in which reactive oxygen species stress was associated with static magnetic field exposure in *E. coli* exposed to a 250 mT static magnetic field, an effect measured using chemical fluorescent probes, electron paramagnetic resonance spectroscopy, and a genetically engineered redox biosensor [8]. This analysis revealed a transcription response similar to H_2_O_2_ exposure.

Compared to the work presented here, however, all the studies have one thing in common: *E. coli* gene expression is studied under hypermagnetic conditions, namely, under fields added on top of the geomagnetic field, which is approximately 50 *µ*T. As such, even the smallest magnetic fields investigated (at 1.1 mT in the Cairo *et al.* study) were still 22-fold stronger than normal geomagnetic conditions, with exposures such as Li *et al.*’s 250 mT static magnetic field at a massive 5,000-fold stronger than the geomagnetic field *E. coli* has evolved under. Conversely, the results presented here investigate the effects on transcriptional gene expression of *E. coli* under hypomagnetic conditions, that is, under a total field strength below that of geomagnetic exposure. This was achieved by growing *E. coli* in a hypomagnetic chamber that shielded the cultures from the bulk of the geomagnetic field to a total field strength of approximately 19 nT. This was achieved by growing *E. coli* in a hypomagnetic chamber that shielded the cultures from the bulk of the geomagnetic field to a total field strength of approximately 19 nT. This difference in field strength can be thought of as large and small in different ways: it is a 2600-fold reduction in field strength, but it is also only a 50 *µ*T difference in absolute terms.

Another aspect of the results presented here is that we investigate lag phase. The other *E. coli* gene expression studies examined cells at high titre in late growth phase or stationary phase, or, in one case, colonies on solid media, which represent cells at mixed stages of growth. Conversely, guided by our prior work showing that hypomagnetic exposure causes *E. coli* to differ in lag phase [1], we chose to sample RNA at 90 minutes post-inoculation. This time was chosen because it is the point where the optical density difference between hypomagnetic and geomagnetic *E. coli* growth curves is most pronounced, with the hypomagnetic culture’s optical density typically about 60% of the geomagnetic culture’s.

## 2. Materials and Methods

### 2.1. Cells and Culture Conditions

As described in our prior work [1], *E. coli* K-12 (ATCC 29425) colonies were isolated on LB agar plates and grown in unsupplemented liquid LB medium (Lennox formulation; pH = 7.0) at 37 *^◦^*C under ambient CO_2_ concentrations and agitated at 100 cycles per minute. Liquid cultures were grown in 25 mL volumes in 50 mL opaque black conical tubes.

Cultures for RNA extraction were grown from seed cultures. A colony was picked two days prior to RNA extraction and grown in LB medium under geomagnetic conditions for 24 hours. One day prior to RNA extraction, 1 mL of that culture was inoculated into 25 mL of fresh LB and placed in either hypomagnetic or geomagnetic conditions, and allowed to grow for 24 hours. Each culture for RNA extraction was 1 mL of a seed culture added to 25 mL of fresh LB and then grown under either hypomagnetic or geomagnetic conditions for 90 minutes. There are four possible RNA extraction cultures, each abbreviated by its seed-culture condition followed by its 90-minute exposure condition: HH (hypomagnetic seed, hypomagnetic exposure); HG (hypomagnetic seed, geomagnetic exposure); GG (geomagnetic seed, geomagnetic exposure); and GH (geomagnetic seed, hypomagnetic exposure). Inoculations were unavoidably performed in a biohood under geomagnetic conditions, but this step was kept as brief as possible (approximately 2 minutes) to minimize exposure to geomagnetic conditions.

### 2.2. Experimental Design

To identify the gene-expression changes underlying the lengthened lag phase caused by hypomagnetic exposure, transcriptomes were compared between *E. coli* grown under hypomagnetic and geomagnetic conditions. Sampling was performed 90 minutes after inoculation, the time at which the optical density of the two conditions differs most in the growth curves reported previously [1].

Experiment A was an initial comparison of two cultures inoculated from a single geomagnetically grown seed, one returned to geomagnetic conditions and one moved to hypomagnetic conditions (GG and GH, respectively). Because both cultures shared a geomagnetic seed, genes differentially expressed in this comparison could reflect either a response to the hypomagnetic condition itself, or a response to the change from geomagnetic to hypomagnetic conditions. Experiment B was performed to distinguish these possibilities. It added seed cultures grown under hypomagnetic conditions, yielding all four seed-condition followed by 90-minute exposure-condition combinations (GG, GH, HG, and HH). This design ensured that the effects of the magnetic field conditions are being compared, rather than the effects of changing magnetic field conditions.

### 2.3. Hypomagnetic Environment

As in prior work [1], a Twinleaf MS-1L hypomagnetic chamber installed inside an agitating incubator provided the hypomagnetic conditions used to test *E. coli*. The chamber was degaussed and measured to have an internal magnetic field strength of 19.9 nT. No active field compensation was used in these experiments. The residual field strength was measured with a QuSpin sensor using QuSpin 2FM UI V7.6.2 software.

### 2.4. RNA Extraction

The 90-minute lag phase cultures were removed from the incubator and immediately spun, in their 50 mL conical tubes, at 5 *^◦^*C and 5,000 rpm for 2 minutes. All medium was drained and cell pellets were resuspended in 100 *µ*L of freshly made 10 mg/mL lysozyme TE solution, which was then transferred to a 2 mL tube and vortexed. Next, 0.5 *µ*L of 10% SDS was added and the tubes were re-vortexed and then allowed to sit at room temperature for 5 minutes, after which they were vortexed for 30 seconds. Then, 350 *µ*L of RLT buffer from the QIAGEN RNeasy Mini Kit [9] was added to each sample, followed by vortexing, and the mixture was loaded onto an Invitrogen PureLink RNA Mini Kit Homogenizer [10] and spun at 12,000 rpm for 2 minutes at 5 *^◦^*C. The flow-through was then processed through the QIAGEN RNeasy Mini Kit starting at step two of the manufacturer-supplied procedure. Total RNA was eluted in RNase-free water.

### 2.5. Transcriptomic Processing

CD Genomics (Shirley, New York) was contracted to perform ribosomal RNA depletion and to construct and sequence transcriptome libraries on an Illumina MiSeq PE150. They also performed initial bioinformatic processing using edgeR [11], calculating fold change and *p*-values. To account for multiple hypothesis testing across all genes, *p*-values were adjusted to a false discovery rate (FDR). Fold change *≥* 2 and FDR *≤* 0.05 were set as screening criteria. Because batch effects commonly dominate transcriptomic datasets [12], genes were further prioritized by reproducibility: a gene was considered reliably differentially expressed only if it met both criteria in at least 4 of the 7 comparisons.

## 3. Results

Differential expression was assessed across seven pairwise comparisons: the single GG vs GH comparison of Experiment A, and the six pairwise comparisons among the four cultures of Experiment B. Across all seven, 343 genes met both screening criteria (fold change *≥* 2, FDR *≤* 0.05) in at least one comparison. Experiment A’s GG vs GH comparison yielded 34 differentially expressed genes; counts for the six Experiment B comparisons are shown in Figure 1 and range from 2 to 242. The complete list of differentially expressed genes for each comparison is provided in the Supplementary Information.

**Fig. 1.**
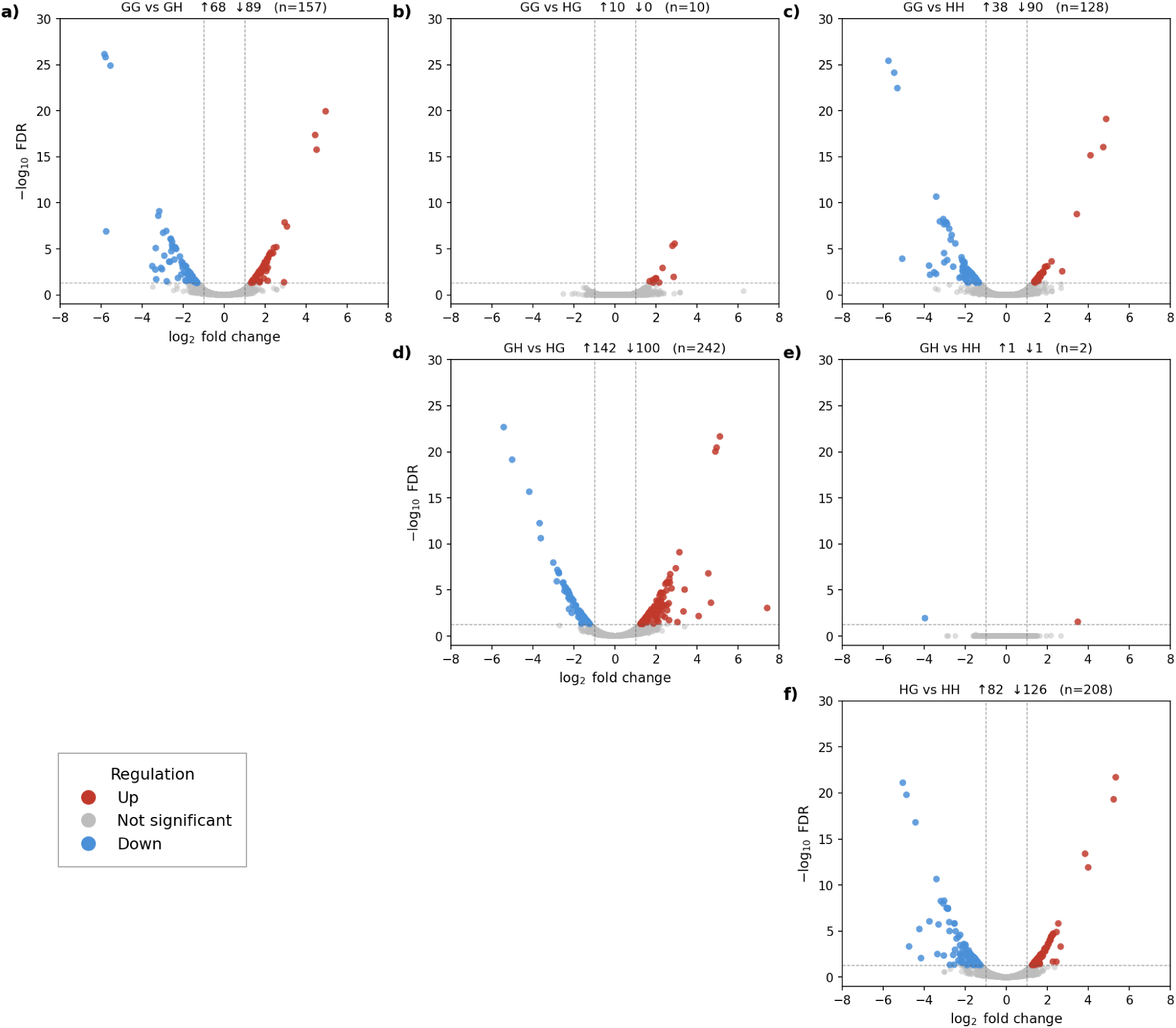
Differential gene expression across the six pairwise comparisons of Experiment B. Each panel shows a volcano plot of one comparison, plotting log_2_ fold change against *−* log_10_ FDR. Genes meeting both screening criteria (FDR *≤* 0.05 and *|* log_2_ fold change*| ≥* 1) are colored by direction of change: red indicates higher expression in the first-listed condition, blue the second; grey indicates genes not meeting both criteria. Dashed lines mark the screening thresholds. Panel headers report the number of up- and down-regulated genes and the total meeting both criteria (n). Conditions are abbreviated by seed-culture condition followed by 90-minute exposure condition (G, geomagnetic; H, hypomagnetic).

Of these genes, 81 met the screening criteria in at least 4 of the 7 comparisons. Among these 81 reproducibly differentially expressed genes, the direction of differential expression tracked the 90-minute exposure condition rather than the seed culture condition: comparisons between cultures differing in their 90-minute field exposure yielded large numbers of differentially expressed genes (128–242), whereas the two comparisons between cultures sharing a 90-minute exposure condition but differing in seed condition (GG vs HG and GH vs HH) yielded few (10 and 2, respectively).

## 4. Discussion

One possible explanation for there being consistently differentially expressed genes between hypomagnetic- and geomagnetic-exposed cultures of *E. coli* is that it reflects nothing more than the two cultures being differentially in lag phase. One could argue that hypomagnetic conditions do not create a distinct metabolic state or regulatory response, but rather trigger the known regulatory program of lag phase. If this were the case, one would expect the lag-phase regulatory program to resemble the 81 genes differentially regulated between hypomagnetic and geomagnetic conditions.

Perhaps the best study describing the lag-phase program is Rolfe *et al.*, which characterizes the lag-phase transcriptome — including 945 genes upregulated within 20 minutes of inoculation — of *Salmonella enterica* serovar Typhimurium, a close relative of *E. coli* [13]. Rolfe *et al.* argue that lag phase is not merely a quiescent delay before logarithmic growth but a distinct growth phase, characterized by induction of transcription and translation machinery, nucleotide synthesis, Fe–S cluster assembly, aerobic respiration, lipid/LPS synthesis, phosphate acquisition, and transient metal accumulation.

There are similarities between the 81 differentially regulated genes under hypomagnetic conditions and those described by Rolfe *et al.* Notably, this includes iron-metabolism genes such as *sufABCDS* and *ynfU*, all of which are upregulated in the hypomagnetic cultures as well as in lag phase. However, the picture concerning iron is not so simple: nearly the entire siderophore iron-scavenging enterobactin pathway (*entABCDEFH*, *fepA*, *flu*, *cirA*, *fhuE*, *fecI*, and *fecR*) is also upregulated under hypomagnetic conditions, which is not typical of lag phase [13]. This suggests that the hypomagnetic conditions cause *E. coli* to perceive and react to iron unavailability.

Similarly, much of the structural flagellar regulon (*flgA*, *fliA*, *fliF*, *fliG*, *fliO*, *fliZ*) in our work is downregulated under hypomagnetic conditions, which is not observed in Rolfe *et al.* Because building flagella is a metabolically expensive task typically associated with cells preparing for growth, this is the opposite of a lag-phase expression pattern; *E. coli* typically shuts down flagellar synthesis as a stress response. This flagellar downregulation is spread across numerous operons and is likely itself a consequence of the downregulation of *fliA* (*σ*^28^), the sigma factor controlling late flagellar gene expression, and *fliZ*, an important positive regulator of the flagellar program. Iron stress is consistent with flagellar-biosynthesis downregulation [14]. Other stress-related genes were also upregulated under hypomagnetic conditions: *ibpA* and *ibpB*, heat-shock proteins that mitigate damage from unfolded proteins.

Another category of differentially regulated genes comprises sugar-metabolism pathways: the glucose PTS (*ptsGHI*), fructose PTS (*fruABK*), mannose PTS (*manXYZ*), and the N-acetylgalactosamine (GalNAc) pathway (*agaSV*). All of these are associated with the phosphotransferase system (PTS), which simultaneously phosphorylates and transports sugars directly into glycolysis in a single step. Coupled with the downregulation of the entire sorbitol pathway (*srlABDER*, *gutM*) — a more biologically costly route — and the downregulation of *ppsA*, which diverts pyruvate toward PEP for gluconeogenesis, we see a selective carbon-metabolism program that privileges fast, easy energy from glycolysis over biosynthetic complexity and growth. This may be associated with the upregulation of *treA*, which is implicated in osmotic protection and stress recovery in carbon-source selection.

Collectively, these differentially regulated genes support several conclusions. First, while some evidence for differential lag phase is present among the differentially expressed genes, there is abundant evidence that they also represent a distinct metabolic state layered on top of lag phase. The strongest evidence for this is the atypical iron stress signaled by the regulation of siderophore and flagellar genes. Combined with the PTS-mediated, glycolysis-privileged metabolic program and numerous stress-response genes, one sees a metabolic state that builds up energy and conserves resources as if uncertain whether to commit to growth. This is consistent with the observed phenotype of a prolonged lag phase.

Most of these differentially expressed genes are likely second- or higher-order effects of the modulation of whatever component of *E. coli* actually senses the hypomagnetic conditions. There is insufficient information in this dataset to conclude what biomolecule the *E. coli* magnetosensor is. However, a hypothesis can be advanced. A leading contender for magnetosensing in biology — where large conductive channels and magnetosomes can be ruled out — is the radical pair mechanism [15]. This mechanism is likely responsible for the magnetic-field-dependent fluorescence of the flavoprotein MagLOV2, which has been measured in live *E. coli* [16]. The redox states of flavins in flavoproteins are one of many redox-sensitive systems *E. coli* uses to monitor its redox state, alongside the NADH/NAD^+^ ratio, Fe^2+^ versus Fe^3+^ balance, Fe–S cluster integrity, proton motive force, and oxygen availability. The hypothesis is that some of this redox monitoring is altered by hypomagnetic conditions, so that cells receive ambiguous redox signals and mount a conservative stress response. While the hypomagnetic response may be a distinct stress response from the hypermagnetic and ROS-stress responses reported in prior high-field studies, this hypothesis is consistent with those results.

Future research suggested by these results includes a time-of-exposure analysis. Just as normal lag phase is a structured metabolic state that passes through a series of steps, the hypomagnetic response likely is as well. Transcriptomic samples taken at earlier points in the growth curve may allow more precise identification of the first-order effects of hypomagnetic exposure, and thus of the magnetosensing biomolecule. Other ′omic layers may further elucidate the genes necessary and sufficient for magnetosensing. Because the hypomagnetic response involves delayed growth, selective conditions might be established to evolve a less-sensitive strain, adding a genomic layer of context. Similarly, because ROS has been implicated in magnetosensitivity, a metabolomic layer may be quite informative.

## Supplemental Information

The complete differential expression tables for all seven pairwise comparisons, including fold-change and FDR values for every gene, are available at Zenodo (https://doi.org/10.5281/zenodo.21709561). Tables SI1 and SI2 list the 81 genes that met the screening criteria in at least four of the seven comparisons, grouped by functional category. Direction of regulation is reported relative to the hypomagnetic 90-minute exposure condition.

**Table SI1:**
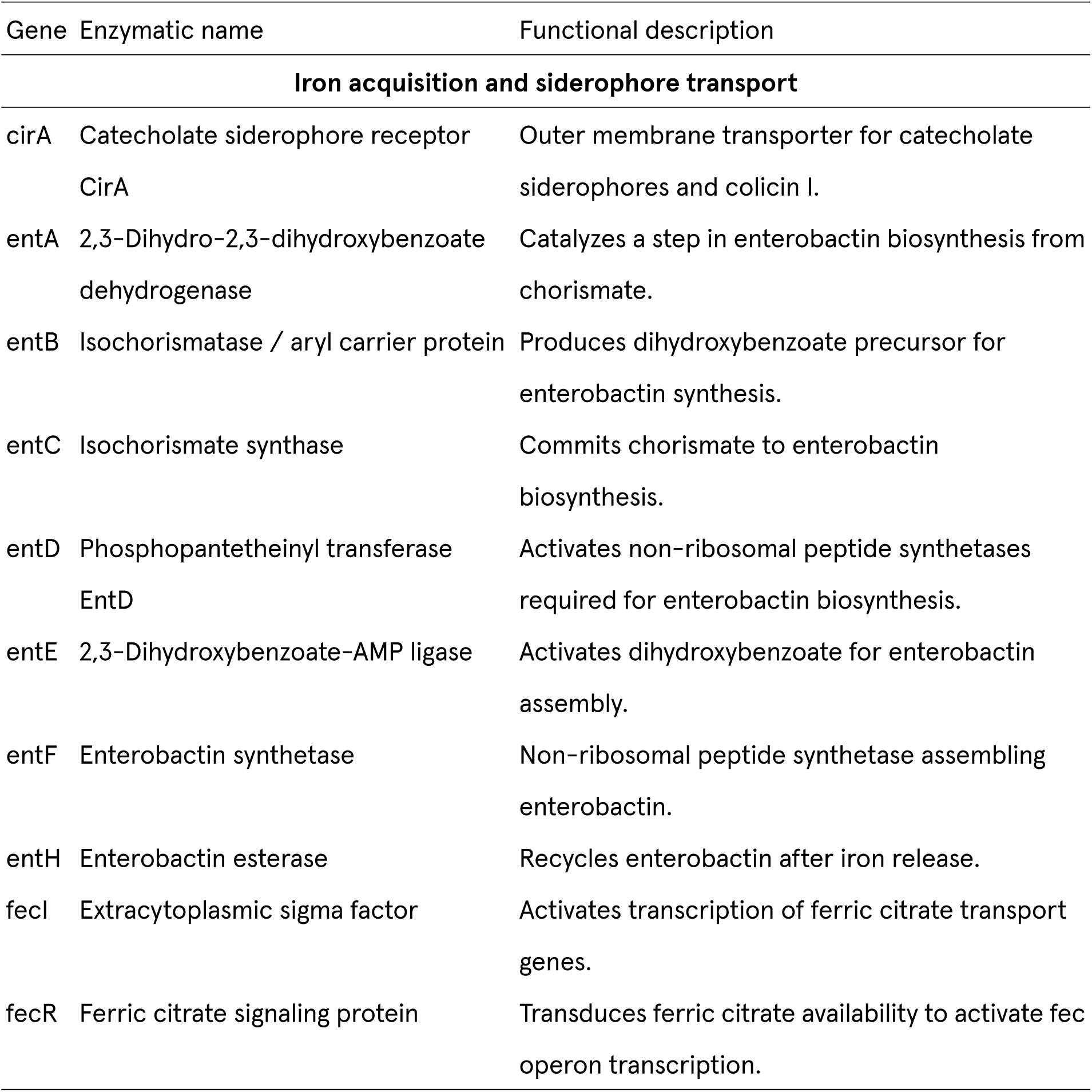
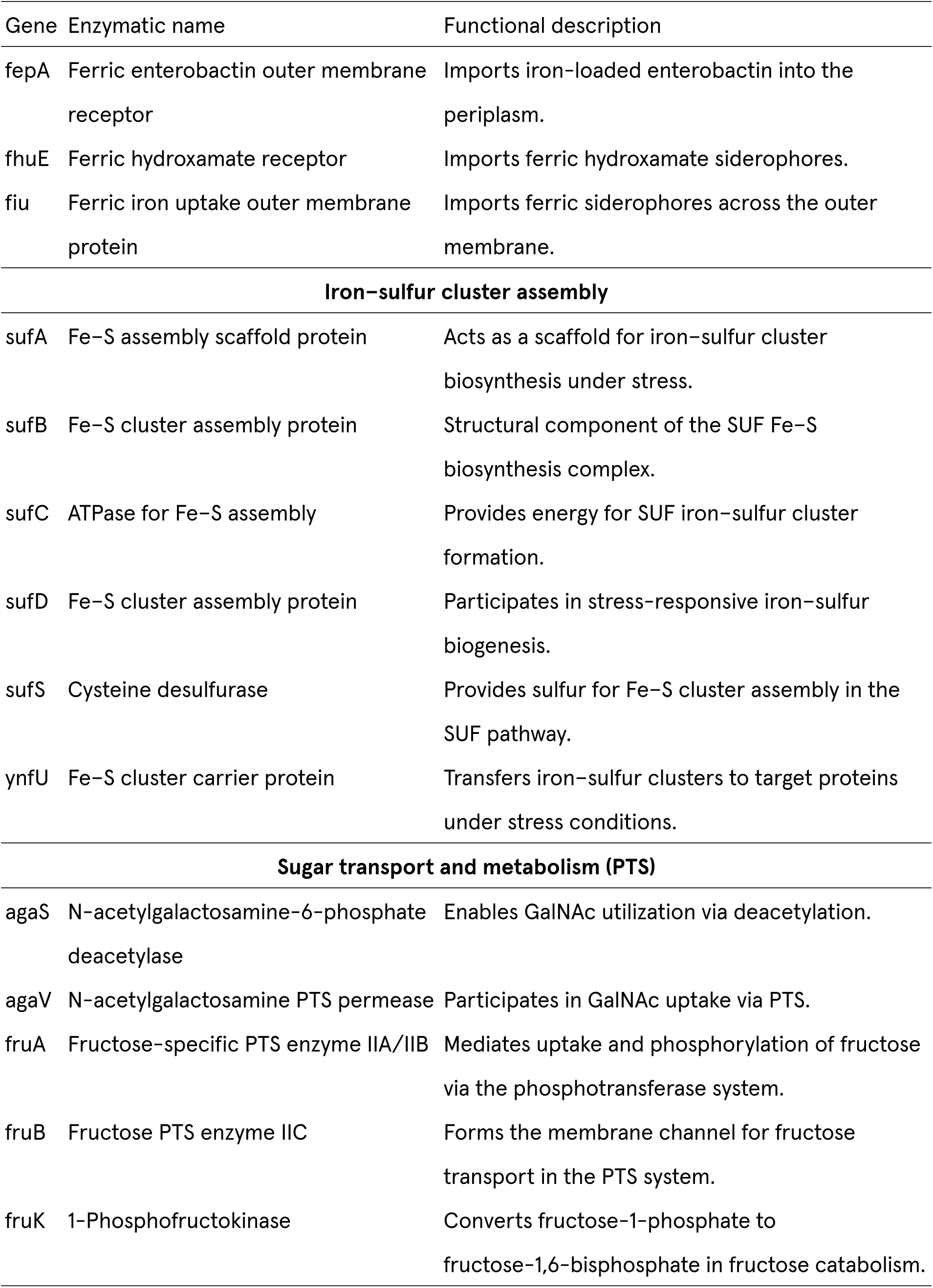
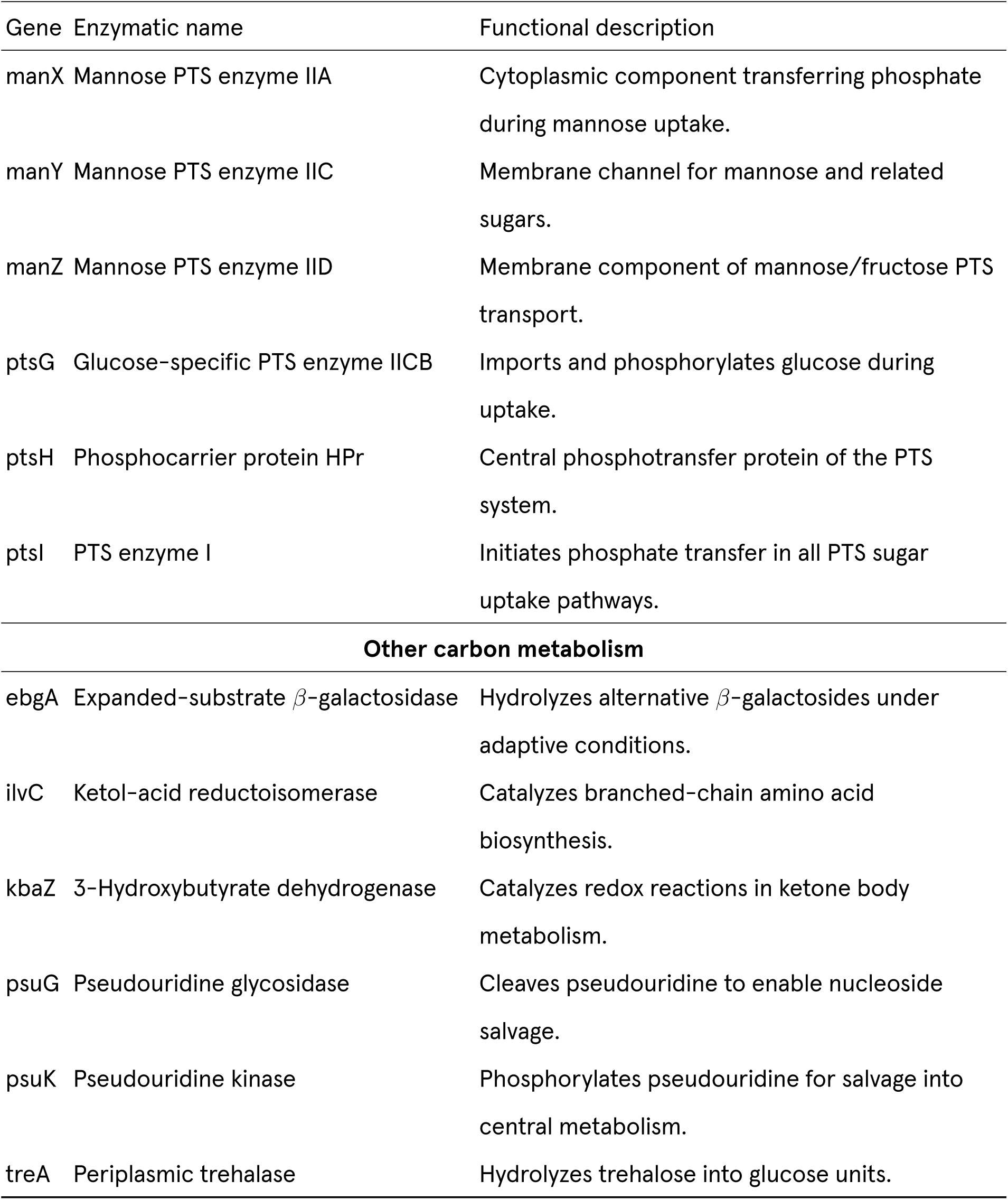
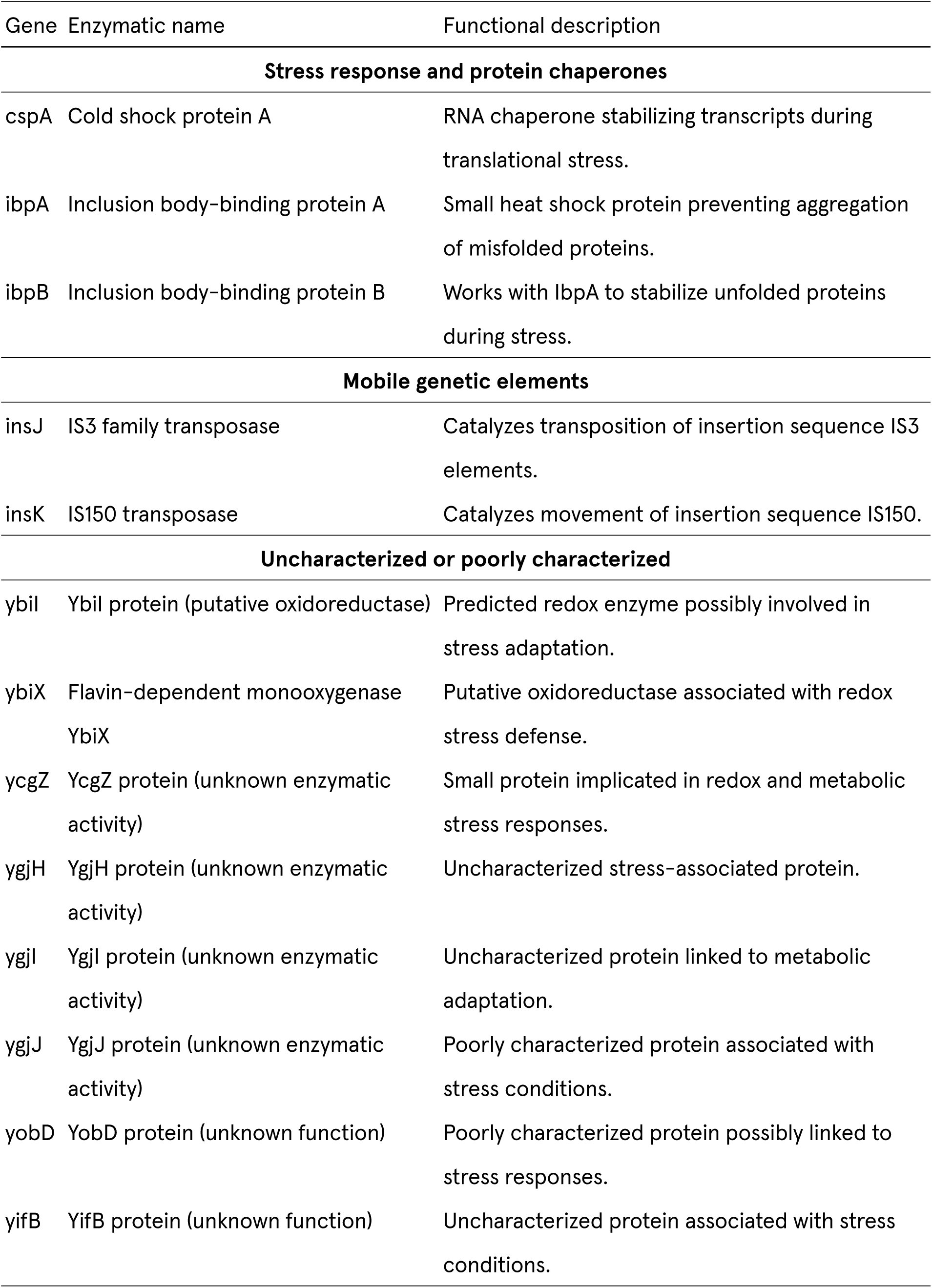
Genes with higher expression under hypomagnetic 90-minute exposure, grouped by functional category.

| Gene | Enzymatic name | Functional description |
| --- | --- | --- |
| **Iron acquisition and siderophore transport** |  |  |
| cirA | Catecholate siderophore receptor<br>CirA | Outer membrane transporter for catecholate siderophores and colicin I. |
| entA | 2,3-Dihydro-2,3-dihydroxybenzoate dehydrogenase | Catalyzes a step in enterobactin biosynthesis from chorismate. |
| entB | Isochorismatase / aryl carrier protein | Produces dihydroxybenzoate precursor for enterobactin synthesis. |
| entC | Isochorismate synthase | Commits chorismate to enterobactin biosynthesis. |
| entD | Phosphopantetheinyl transferase<br>EntD | Activates non-ribosomal peptide synthetases required for enterobactin biosynthesis. |
| entE | 2,3-Dihydroxybenzoate-AMP ligase | Activates dihydroxybenzoate for enterobactin assembly. |
| entF | Enterobactin synthetase | Non-ribosomal peptide synthetase assembling enterobactin. |
| entH | Enterobactin esterase | Recycles enterobactin after iron release. |
| fecI | Extracytoplasmic sigma factor | Activates transcription of ferric citrate transport genes. |
| fecR | Ferric citrate signaling protein | Transduces ferric citrate availability to activate fec operon transcription. |
| fepA | Ferric enterobactin outer membrane receptor | Imports iron-loaded enterobactin into the periplasm. |
| fhuE | Ferric hydroxamate receptor | Imports ferric hydroxamate siderophores. |
| fiu | Ferric iron uptake outer membrane protein | Imports ferric siderophores across the outer membrane. |
| **Iron-sulfur cluster assembly** |  |  |
| sufA | Fe-S assembly scaffold protein | Acts as a scaffold for iron-sulfur cluster biosynthesis under stress. |
| sufB | Fe-S cluster assembly protein | Structural component of the SUF Fe-S biosynthesis complex. |
| sufC | ATPase for Fe-S assembly | Provides energy for SUF iron-sulfur cluster formation. |
| sufD | Fe-S cluster assembly protein | Participates in stress-responsive iron-sulfur biogenesis. |
| sufS | Cysteine desulfurase | Provides sulfur for Fe-S cluster assembly in the SUF pathway. |
| ynfU | Fe-S cluster carrier protein | Transfers iron-sulfur clusters to target proteins under stress conditions. |
| **Sugar transport and metabolism (PTS)** |  |  |
| agaS | N-acetylgalactosamine-6-phosphate deacetylase | Enables GalNAc utilization via deacetylation. |
| agaV | N-acetylgalactosamine PTS permease | Participates in GalNAc uptake via PTS. |
| fruA | Fructose-specific PTS enzyme IIA/IIB | Mediates uptake and phosphorylation of fructose via the phosphotransferase system. |
| fruB | Fructose PTS enzyme IIC | Forms the membrane channel for fructose transport in the PTS system. |
| fruK | 1-Phosphofructokinase | Converts fructose-1-phosphate to fructose-1,6-bisphosphate in fructose catabolism. |
| manX | Mannose PTS enzyme IIA | Cytoplasmic component transferring phosphate during mannose uptake. |
| manY | Mannose PTS enzyme IIC | Membrane channel for mannose and related sugars. |
| manZ | Mannose PTS enzyme IID | Membrane component of mannose/fructose PTS transport. |
| ptsG | Glucose-specific PTS enzyme IICB | Imports and phosphorylates glucose during uptake. |
| ptsH | Phosphocarrier protein HPr | Central phosphotransfer protein of the PTS system. |
| ptsI | PTS enzyme I | Initiates phosphate transfer in all PTS sugar uptake pathways. |
| **Other carbon metabolism** |  |  |
| ebgA | Expanded-substrate β-galactosidase | Hydrolyzes alternative β-galactosides under adaptive conditions. |
| ilvC | Ketol-acid reductoisomerase | Catalyzes branched-chain amino acid biosynthesis. |
| kbaZ | 3-Hydroxybutyrate dehydrogenase | Catalyzes redox reactions in ketone body metabolism. |
| psuG | Pseudouridine glycosidase | Cleaves pseudouridine to enable nucleoside salvage. |
| psuK | Pseudouridine kinase | Phosphorylates pseudouridine for salvage into central metabolism. |
| treA | Periplasmic trehalase | Hydrolyzes trehalose into glucose units. |
| **Stress response and protein chaperones** |  |  |
| cspA | Cold shock protein A | RNA chaperone stabilizing transcripts during translational stress. |
| ibpA | Inclusion body-binding protein A | Small heat shock protein preventing aggregation of misfolded proteins. |
| ibpB | Inclusion body-binding protein B | Works with IbpA to stabilize unfolded proteins during stress. |
| **Mobile genetic elements** |  |  |
| insJ | IS3 family transposase | Catalyzes transposition of insertion sequence IS3 elements. |
| insK | IS150 transposase | Catalyzes movement of insertion sequence IS150. |
| **Uncharacterized or poorly characterized** |  |  |
| ybil | Ybil protein (putative oxidoreductase) | Predicted redox enzyme possibly involved in stress adaptation. |
| ybiX | Flavin-dependent monooxygenase<br>YbiX | Putative oxidoreductase associated with redox stress defense. |
| ycgZ | YcgZ protein (unknown enzymatic activity) | Small protein implicated in redox and metabolic stress responses. |
| ygjH | YgjH protein (unknown enzymatic activity) | Uncharacterized stress-associated protein. |
| ygjI | YgjI protein (unknown enzymatic activity) | Uncharacterized protein linked to metabolic adaptation. |
| ygjJ | YgjJ protein (unknown enzymatic activity) | Poorly characterized protein associated with stress conditions. |
| yobD | YobD protein (unknown function) | Poorly characterized protein possibly linked to stress responses. |
| yifB | YifB protein (unknown function) | Uncharacterized protein associated with stress conditions. |

**Table SI2:**
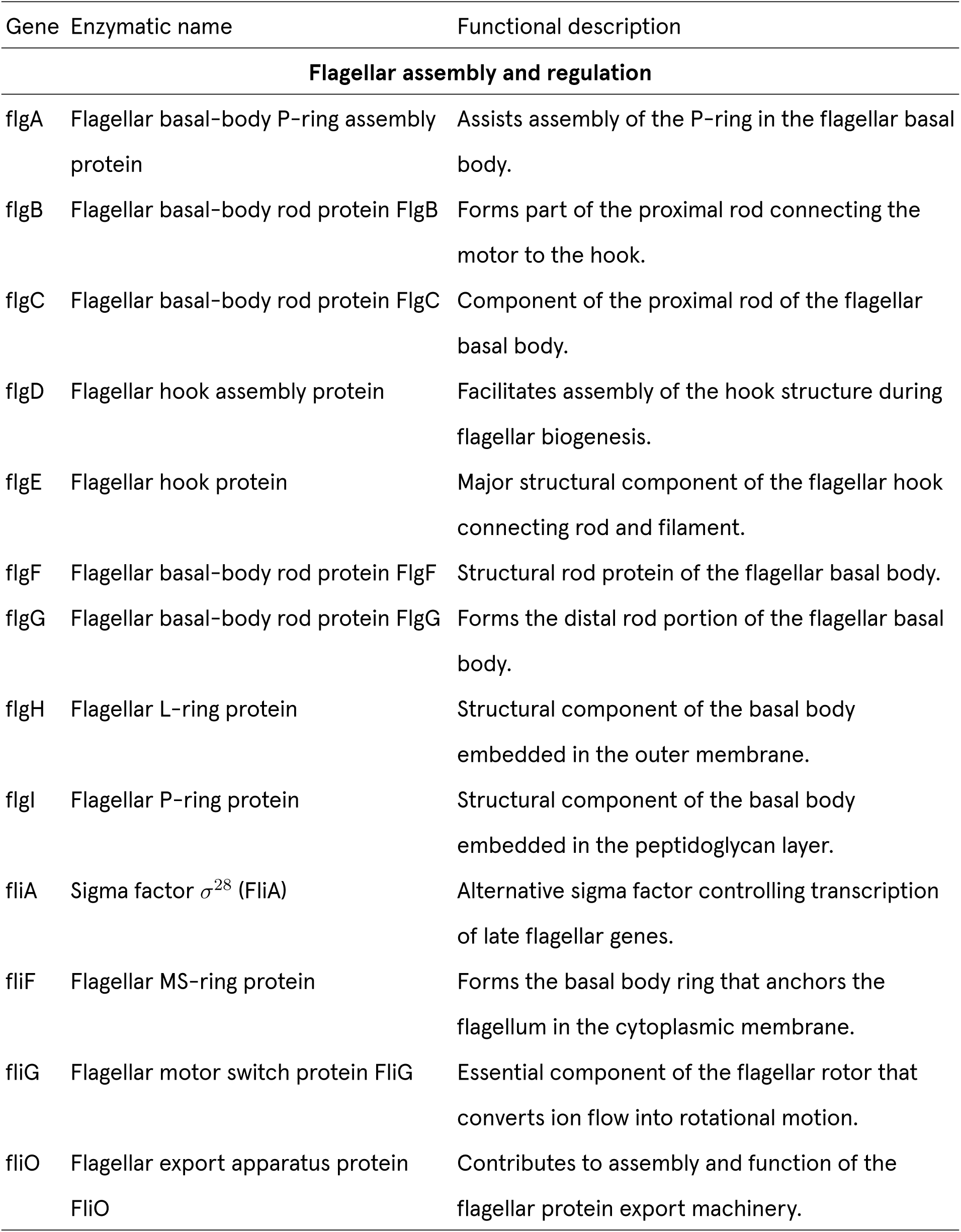
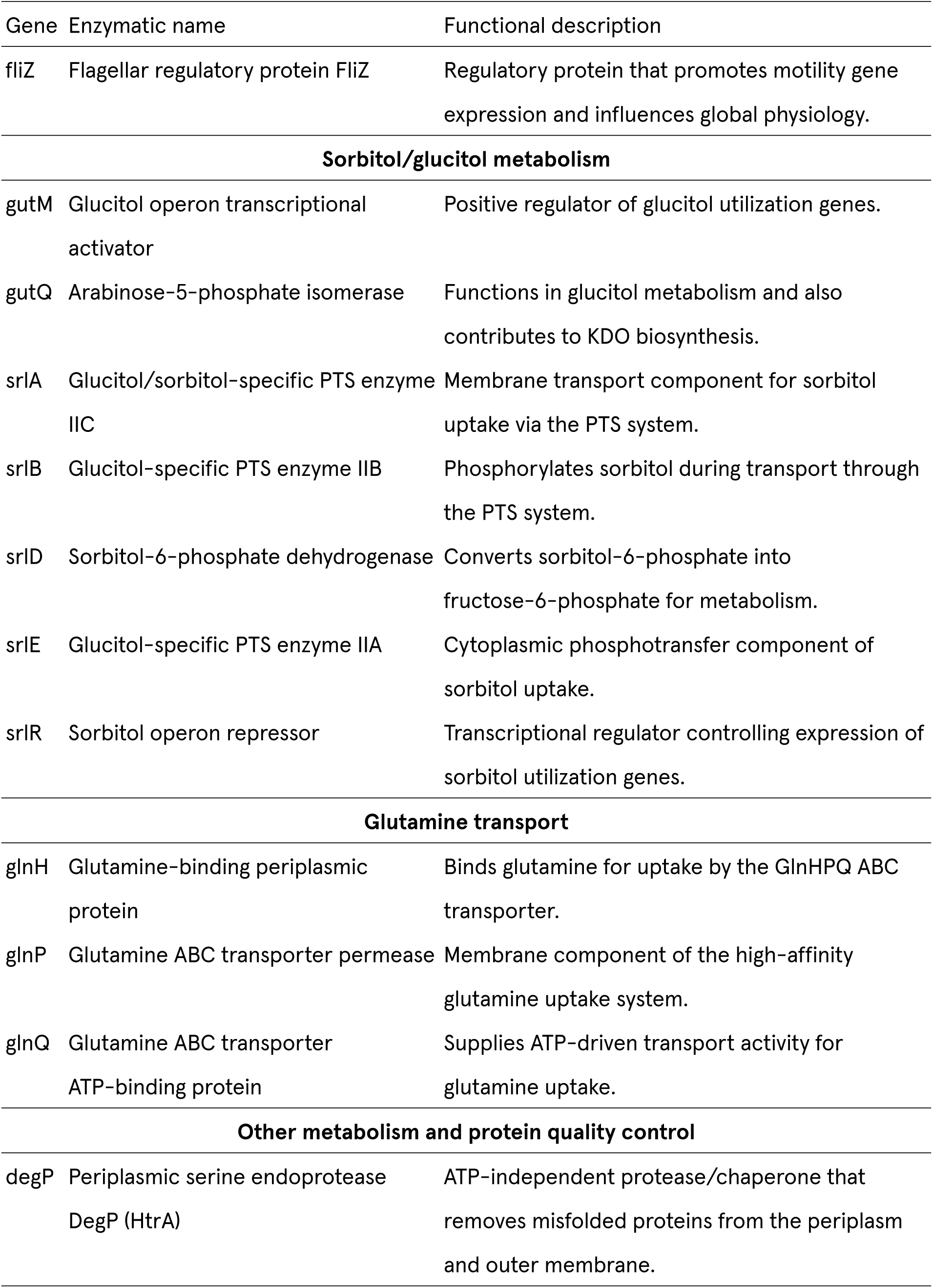
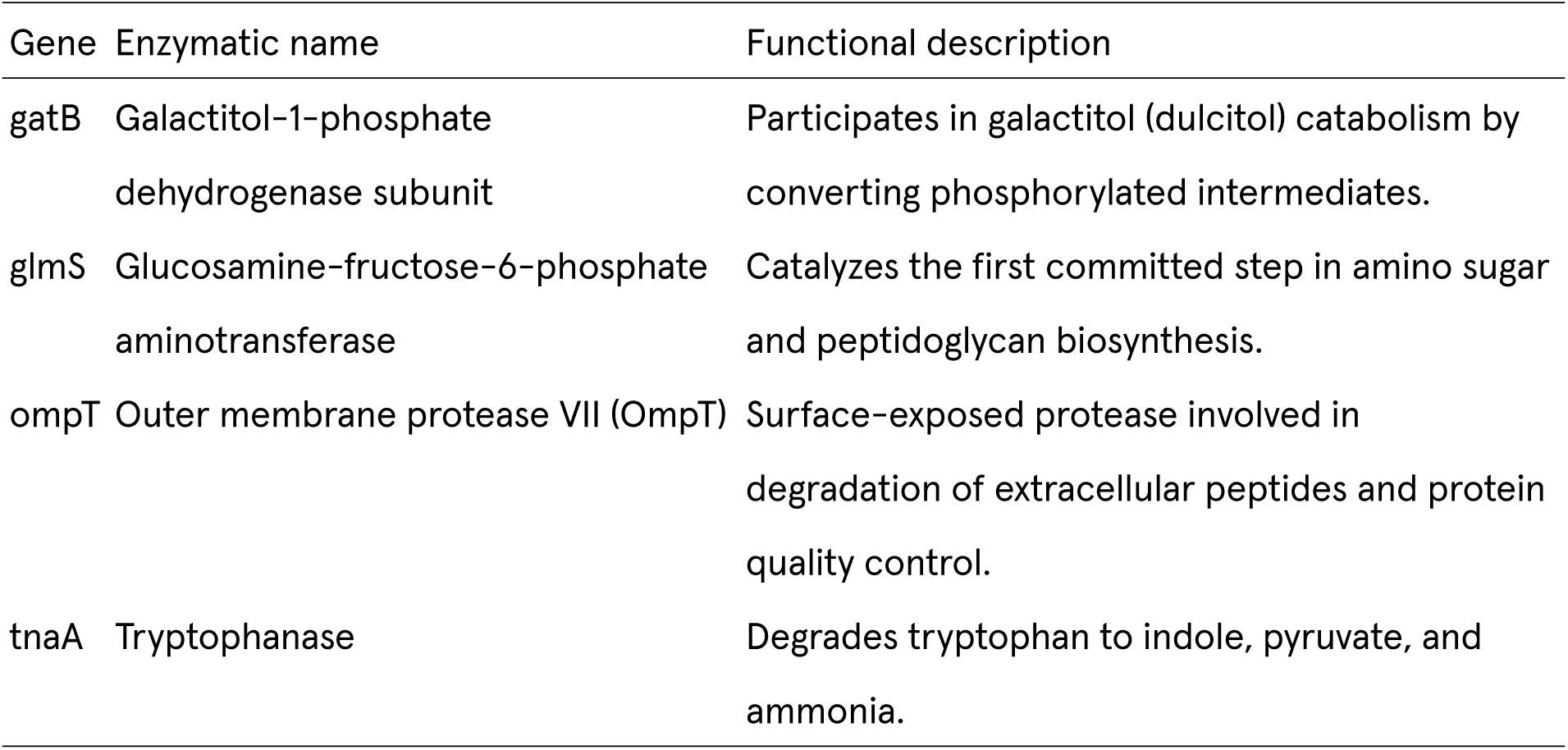
Genes with lower expression under hypomagnetic 90-minute exposure, grouped by functional category.

## 8. Author Contributions

Following the CRediT taxonomy [17]:

1. 1) Michael Montague: conceptualization; data curation; formal analysis; investigation; methodology; software; validation; visualization; and writing (original draft)
2. 2) Morgan L. Sosa: data curation; formal analysis; software; visualization; and writing (review and editing)
3. 3) Clarice D. Aiello: conceptualization; funding acquisition; validation; and writing (review and editing)

The Quantum Biology Institute is a California non-profit 501(c)(3) focused research organization that performs basic research underpinning the quantum biology field in an open-science fashion.

## References

[1] M. Montague, A. Lodesani, and C. D. Aiello, Escherichia coli k12 exhibits a ∼50% longer lag phase, but no difference in log phase growth rate, under hypomagnetic conditions (19 nt) 10.64898/2026.04.13.717819 (2026).

[2] P. Cairo, B. Greenebaum, and E. Goodman, Magnetic field exposure enhances mrna expression of *σ*^32^ in *e. coli*, Journal of Cellular Biochemistry 68, 1 (1998).

[3] L. Potenza, L. Ubaldi, R. De Sanctis, R. De Bellis, L. Cucchiarini, and M. Dachà, Effects of a static magnetic field on cell growth and gene expression in *escherichia coli*, Mutation Research/Genetic Toxicology and Environmental Mutagenesis 561, 53 (2004).

[4] E. Twiss, A. M. Coros, N. P. Tavakoli, and K. M. Derbyshire, Transposition is modulated by a diverse set of host factors in *escherichia coli* and is stimulated by nutritional stress, Molecular Microbiology 57, 1593 (2005).

[5] K. Tsuchiya, K. Okuno, T. Ano, K. Tanaka, H. Takahashi, and M. Shoda, High magnetic field enhances stationary phase-specific transcription activity of *escherichia coli*, Bioelectrochemistry and Bioenergetics 48, 383 (1999).

[6] E. M. Goodman, B. Greenebaum, and M. T. Marron, Magnetic fields alter translation in *escherichia coli*, Bioelectromagnetics 15, 77 (1994).

[7] S. G. Huwiler, C. Beyer, J. Fröhlich, H. Hennecke, T. Egli, D. Schürmann, H. Rehrauer, and H.-M. Fischer, Genome-wide transcription analysis of *escherichia coli* in response to extremely low-frequency magnetic fields, Bioelectromagnetics 33, 488 (2012).

[8] H. Li, Y. Fang, and J. Huang, Reactive oxygen species mediate bioeffects of static magnetic field via impairment of long-chain fatty acid degradation in *escherichia coli*, Frontiers in Microbiology 16, 1586233 (2025).

[9] QIAGEN, RNeasy kits, https://www.qiagen.com/us/products/discovery-and-translational-research/dna-rna-purification/rna-purification/total-rna/rneasy-kits, accessed: 2026-07-29.

[10] Thermo Fisher Scientific, PureLink™ RNA mini kit, https://www.thermofisher.com/order/catalog/product/12183026, accessed: 2026-07-29.

[11] M. D. Robinson, D. J. McCarthy, and G. K. Smyth, edgeR: a Bioconductor package for differential expression analysis of digital gene expression data, Bioinformatics 26, 139 (2010).

[12] Y. Yu, Y. Mai, Y. Zheng, and L. Shi, Assessing and mitigating batch effects in large-scale omics studies, Genome Biology 25, 254 (2024).

[13] M. D. Rolfe, C. J. Rice, S. Lucchini, C. Pin, A. Thompson, A. D. S. Cameron, M. Alston, M. F. Stringer, R. P. Betts, J. Baranyi, M. W. Peck, and J. C. D. Hinton, Lag phase is a distinct growth phase that prepares bacteria for exponential growth and involves transient metal accumulation, Journal of Bacteriology 194, 686 (2012).

[14] J. P. McHugh, F. Rodríguez-Quiñones, H. Abdul-Tehrani, D. A. Svistunenko, R. K. Poole, C. E. Cooper, and S. C. Andrews, Global iron-dependent gene regulation in *escherichia coli*: a new mechanism for iron homeostasis, Journal of Biological Chemistry 278, 29478 (2003).

[15] C. D. Aiello, B. L. Ross, A. Lodesani, and M. L. Sosa, A physicist-friendly primer on the Hamiltonian for quantum sensing in proteins: analytical expressions and insights for a toy model of the radical-pair mechanism (2026), arXiv:2604.18608 [physics.bio-ph].

[16] B. L. Ross, A. Lodesani, and C. D. Aiello, The magnetic field-dependent fluorescence of MagLOV2 in live bacterial cells is consistent with the radical pair mechanism, bioRxiv 10.64898/2026.02.18.706690 (2026), preprint.

[17] CRediT – Contributor Roles Taxonomy, https://credit.niso.org/, accessed: 2026.

